# Improved high-dimensional multivariate autoregressive model estimation of human electrophysiological data using fMRI priors

**DOI:** 10.1101/2022.11.18.516669

**Authors:** Alliot Nagle, Josh P. Gerrelts, Bryan M. Krause, Aaron D. Boes, Joel E. Bruss, Kirill V. Nourski, Matthew I. Banks, Barry Van Veen

**Affiliations:** Department of Electrical and Computer Engineering, University of Wisconsin, Madison, 53706, WI, USA; Department of Anesthesiology, University of Wisconsin, Madison, 53706, WI, USA; Department Neurology, The University of Iowa, Iowa City, 52242, IA, USA; Department Neurosurgery, The University of Iowa, Iowa City, 52242, IA, USA; Iowa Neuroscience Institute, The University of Iowa, Iowa City, 52242, IA, USA

## Abstract

Multivariate autoregressive (MVAR) model estimation enables assessment of causal interactions in brain networks. However, accurately estimating MVAR models for high-dimensional electrophysiological recordings is challenging due to the extensive data requirements. Hence, the applicability of MVAR models for study of brain behavior over hundreds of recording sites has been very limited. Prior work has focused on different strategies for selecting a subset of important MVAR coefficients in the model and is motivated by the potential of MVAR models and the data requirements of conventional least-squares estimation algorithms. Here we propose incorporating prior information, such as fMRI, into MVAR model estimation using a weighted group LASSO regularization strategy. The proposed approach is shown to reduce data requirements by a factor of two relative to the recently proposed group LASSO method of Endemann et al. (2022) while resulting in models that are both more parsimonious and have higher fidelity to the ground truth. The effectiveness of the method is demonstrated using simulation studies of physiologically realistic MVAR models derived from iEEG data. The robustness of the approach to deviations between the conditions under which the prior information and iEEG data is obtained is illustrated using models from data collected in different sleep stages. This approach will allow accurate effective connectivity analyses over short time scales, facilitating investigations of causal interactions in the brain underlying perception and cognition during rapid transitions in behavioral state.

## 1. Introduction

Directed connectivity and causality have emerged as key attributes for understanding the network interactions underlying cognition and perception. Linear multivariable autoregressive (MVAR) models are the simplest causal model for electrophysiological brain dynamics and have proven remarkably effective at capturing attributes of brain networks. However, MVAR model estimation for high-dimensional networks is notoriously difficult due to the large number of parameters that need to be estimated. Large amounts of data are required to obtain stable parameter estimates for high-dimensional network models, which greatly limits their application. MVAR models assume the data is stationary, an assumption likely to be violated over longer time spans. These large data requirements also preclude dynamic analyses of connectivity critical to understanding cognition and perception in typical environments. For example, Schlogl and Supp (2006) have suggested exceeding 10*MP* samples in time, where *M* is the number of channels and *P* is the model memory, when using ordinary least-squares (OLS) to estimate the model parameters. Studies by Antonacci et al. (2020) and Antonacci et al. (2021) that compare OLS to variations on the Least Absolute Shrinkage and Selection Operator (LASSO) of Tibshirani (1996) confirm that OLS requires 10*MP* to 20*MP* time samples for accurate estimation of model parameters.

Human intracranial electroencephalographic (iEEG) recordings offer a rare opportunity to investigate brain activity at high temporal and spatial resolution. Simultaneous data collection from >200 recording sites is not uncommon in iEEG, allowing for sampling from large areas of the brain involved in sensorimotor processing, memory access and retrieval, executive function, and other processes fundamental to human conscious experience. Multivariate autoregressive (MVAR) models fit to resting state (RS) data can yield detailed estimates of directional connectivity from RS data. However, estimating MVAR models using OLS for networks involving several hundred recording sites requires excessively long data records and motivates development of more advanced algorithmic approaches.

The LASSO approach regularizes the least-squares problem with a penalty on the *ℓ*_1_ norm of the MVAR model coefficients to reduce the amount of data required. Other approaches for reducing the amount of required data include selecting a subset of important MVAR coefficients for inclusion in the model (Marinazzo et al. (2012)) based on mutual information, and restricting the number of past values of each variable using a Bayesian Information Criterion based sequential selection method (Siggiridou and Kugiumtzis (2016)). Endemann et al. (2022) showed recently that regularization using group LASSO (gLASSO) (Yuan and Lin (2006)) dramatically improved model estimates, reducing data requirements by as much as a factor of 10 compared to OLS. The gLASSO regularizes the least-squares problem with the *ℓ*_1_ norm of the *ℓ*_2_ norm of the MVAR coefficients relating the signals at each pair of electrodes. Hence, while the LASSO penalty encourages solutions to the leastsquares problem that are sparse in the number of MVAR coefficients, the gLASSO penalty encourages solutions that are sparse in the number of connections in the MVAR network model. Sparsely connected MVAR models are consistent with hypothesized small-world organization of the brain (Bullmore and Sporns (2009)).

An additional approach for reducing the data requirement of MVAR model estimation is to incorporate prior information about the network into the estimation process. Activity and connectivity profiles from magnetic resonance imaging (MRI) have been used previously to inform analyses of electrophysiological data, for example in EEG source localization (Henson et al. (2010), Lei et al. (2015), Glomb et al. (2020)). Structural and functional connectivity derived from MRI are predictive of connectivity measured electrophysiologically (Hacker et al. (2017)), and thus could prove informative in iEEG network model estimation. Several previous studies have used priors derived from structural or functional brain connectivity measurements to inform functional models in different contexts. For example, Zhu et al. (2013) use structural priors obtained from diffusion tensor imaging (DTI) to differentially penalize the weights in a regression model for resting-state networks based on functional magnetic resonance imaging (fMRI). Similarly, Crimi et al. (2021) construct an MVAR model for fMRI networks where connections are turned on or off based on structural information. Other studies have derived priors from functional connectivity. For example, Yu et al. (2017) use Pearson correlations from fMRI to penalize connections in a group LASSO multiple-region regression for fMRI networks. Brown et al. (2019) estimate sub-networks in the brain that are predictive of outcomes or disease states in a sparse feature selection framework that incorporates priors derived from DTI or fMRI. Similar ideas have also been applied in applications beyond brain networks, such as estimation of MVAR gene networks using prior internal connectivity structure (Charbonnier et al. (2010)). Because the gLASSO operates at the connection level, the gLASSO penalty can be modified to differentially penalize connections based on prior information, an approach called weighted group LASSO (wgLASSO) (Hirose and Konishi (2012)).

In this paper we develop a method for using prior information from fMRI functional connectivity for the estimation of MVAR networks from high-dimensional electrophysiological recordings in order to further reduce data requirements. Neurosurgical patients undergo extensive pre-operative evaluation including MRI. Hence, additional information about the composition of cortical networks in these subjects is readily available and can be exploited to improve the quality of model estimates and further reduce data requirements. Our wgLASSO approach extends the gLASSO method of Endemann et al. (2022) by using this prior information to penalize groups of MVAR coefficients inversely proportional to the likelihood of a connection between the corresponding nodes. To the best of our knowledge, prior information has not previously been used in estimation of MVAR models from electrophysiological data. As in Brown et al. (2019), we use Pearson correlations from the subject’s fMRI BOLD signals to inform the electrophysiological network. The effectiveness of our approach is demonstrated with using simulated data from physiologically plausible iEEG networks derived from resting state human recordings (Endemann et al. (2022)). We also analyze data recorded during overnight sleep to demonstrate that the priors generalize beyond the specific brain state in which they were obtained. The proposed wgLASSO approach substantially reduces data requirements for accurate model estimation, enabling high dimensional network modeling at physiologically meaningful time scales. Furthermore, with relatively long data records the wgLASSO produces sparser models than the gLASSO for comparable data fitting accuracy, suggesting the wgLASSO is estimating models that better fit the small-world organization of the brain. Finally, the proposed wgLASSO approach is also less affected by shrinkage than the gLASSO, which reduces the need for additional debiasing following model selection.

The Methods section describes the iEEG and fMRI data as well as the wgLASSO approach and the performance metrics used to characterize the quality of the estimated models. The performance of the wgLASSO method and the gLASSO approach of Endemann et al. (2022) are reported as a function of simulated model data length in the Results section for both resting state conditions and in different sleep stages. The Discussion section contains a thorough analysis of the relative performance benefits of the proposed wgLASSO method.

## 2. Methods

### 2.1. Participants

Ground-truth networks were derived from resting state iEEG data. Daytime wake data were obtained from 9 neurosurgical patients (4 female, ages 19 - 59 years old; Table 1). Three of these participants were also recorded during overnight sleep, as described previously (Banks et al. (2020)). The patients had been diagnosed with medically refractory epilepsy and were undergoing chronic invasive iEEG monitoring to identify potentially resectable seizure foci. All human participant studies were carried out in accordance with The Code of Ethics of the World Medical Association (Declaration of Helsinki) for experiments involving humans. The research protocols were approved by the University of Iowa Institutional Review Board and the National Institutes of Health. Written informed consent was obtained from all participants. Research participation did not interfere with acquisition of clinically required data. Participants could rescind consent at any time without interrupting their clinical evaluation.

**Table 1:** Participant demographics. Datasets: R = resting state; S = sleep.

| Participant | Age | Gender | $n_{sites}$ | Dataset |
| --- | --- | --- | --- | --- |
| R369 | 30 | M | 216 | R |
| L372 | 34 | M | 178 | R |
| R376 | 48 | F | 207 | R, S |
| L400 | 59 | F | 149 | R |
| L405 | 19 | M | 124 | R |
| L409 | 31 | F | 160 | R |
| R418 | 25 | F | 83 | R, S |
| L423 | 51 | M | 153 | R, S |
| R456 | 31 | M | 195 | R |

### 2.2. Pre-implantation neuroimaging

All participants underwent whole-brain high-resolution T1-weighted structural MRI scans before electrode implantation to aid with electrode localization after surgery. In addition, resting state (RS) fMRI data were collected and used as functional connectivity priors for MVAR model estimation. The scanner was a 3T GE Discovery MR750W with a 32-channel head coil. The pre-electrode implantation anatomical T1 scan (3D FSPGR BRAVO sequence) was obtained with the following parameters: FOV = 25.6 cm, flip angle = 12 deg., TR = 8.50 ms, TE = 3.29 ms, inversion time = 450 ms, voxel size = 1.0 × 1.0 × 0.8 mm. For RS-fMRI, 5 blocks of 5-minute gradient-echo EPI runs (650 volumes) were collected with the following parameters: FOV = 22.0 cm, TR = 2260 ms, TE = 30 ms, flip angle = 80 deg., voxel size = 3.45 × 3.45 × 4.0 mm. For each participant, RS-fMRI runs were acquired in the same session but non-contiguously (dispersed within an imaging session to avoid habituation). Participants were asked to keep their eyes open, and a fixation cross was presented through a projector.

### 2.3. iEEG Recordings

Electrophysiological recordings were obtained using subdural electrode arrays (platinum-iridium discs 2.3 mm diameter, 5-10 mm inter-electrode distance, embedded in a silicon membrane) and depth arrays (8-12 electrodes, 5 mm inter-electrode distance) (Ad-Tech Medical, Oak Creek, WI) (Fig. 1, Supplementary Fig. 2). After rejecting electrodes that were located in seizure foci, white matter, or outside the brain, or for noise reasons (see below), the median number of recording sites across the 9 subjects was 160 (range 83 – 216; Table 1), providing extensive coverage of temporal, frontal, and parietal cortex. A subgaleal electrode, placed over the cranial vertex near midline, was used as a reference in all subjects. All electrodes were placed solely on the basis of clinical requirements, as determined by the team of epileptologists and neurosurgeons Nourski and Howard (2015).

**Figure 1:**
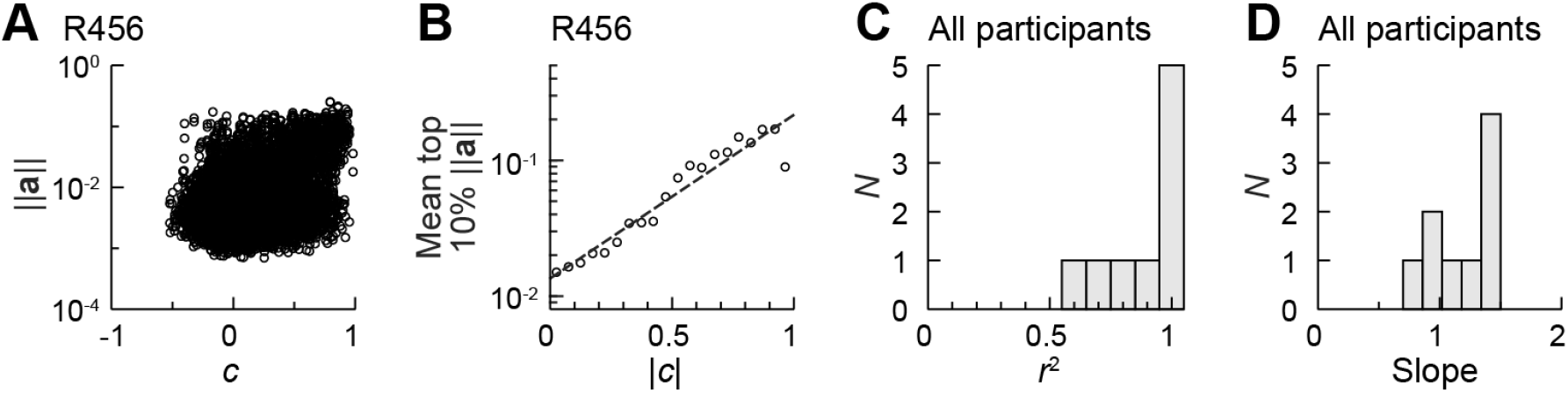
Logarithmic relationship between coefficient norms and fMRI connectivity. (A) Norms of ground truth coefficients as a function of corresponding fMRI Pearson Correlation (PC) values. (B) The data in A were binned according to the magnitude of the fMRI PC values, and the mean upper decile of coefficient norms plotted versus the mean abs(fMRI PC) in each bin. The regression line has slope = 1.20 and *r*^2^ = 0.922. (C, D) Regression lines for all participants were fit as in B. (C) *r*^2^ values for all nine participants. (D) Slopes for all nine participants.

For daytime recordings, no-task, RS data were recorded in the dedicated, electrically shielded suite in The University of Iowa Clinical Research Unit while the participants lay in the hospital bed. Participants typically had their eyes open during data collection. Resting state data were collected a median of 4.8 days (range 1.9 – 21 days) after electrode implantation surgery. Data acquisition was performed using a Neuralynx Atlas System (Neuralynx Inc., Bozeman, MT). Recorded data were amplified, filtered (0.1–500 Hz bandpass, 5 dB/octave rolloff for TDT-recorded data; 0.7–800 Hz bandpass, 12 dB/octave rolloff for Neuralynx-recorded data) and digitized at a sampling rate of 2034.5 Hz (TDT) or 2000 Hz (Neuralynx). The duration of recordings in all participants was at least 6 minutes.

Overnight sleep RS iEEG, scalp EEG, and video data were collected in the dedicated, electrically shielded suite in The University of Iowa Clinical Research Unit while the participants lay in the hospital bed. Sleep data were collected 7 or 8 days after electrode implantation surgery using the same recording system and parameters as for daytime RS data. Stages of sleep were defined manually as described previously (Banks et al. (2020). At least 6 minutes of data were recorded for each sleep stage in each participant.

### 2.4. fMRI Data Analysis

Standard preprocessing was applied to the RS-fMRI data acquired in the pre-implantation scan using FSL’s FEAT pipeline, including spatial alignment and nuisance regression. White matter, cerebrospinal fluid and global ROIs were created using deep white matter, lateral ventricles and a whole brain mask, respectively. Regression was performed using the time series of these three nuisance ROIs as well as 6 motion parameters (3 rotations and 3 translations) and their derivatives, detrended with second order polynomials. Temporal bandpass filtering was 0.008–0.08 Hz. Spatial smoothing was applied with a Gaussian kernel (6 mm full-width at half maximum). The first two images from each run were discarded. Frame censoring was applied when the Euclidean norm of derivatives of motion parameters exceeded 0.5 mm (Power et al. (2012)). All runs were processed in native EPI space, then the residual data were transformed to MNI152 and concatenated.

Functional connectivity was calculated as Pearson correlation coefficients between BOLD signals averaged within voxel groupings. Voxel groupings were based on iEEG electrode location in participant space, with voxels chosen to represent comparable regions of the brain recorded by iEEG electrodes. For each electrode, the anatomical coordinates of the recording site were mapped to the closest valid MRI voxel, E, and a sphere with volume 25 mm^3^ centered on E used as the corresponding recording site. A single time series was computed as the average of the fMRI BOLD signal in these 25 voxels, and a connectivity matrix computed as the Pearson correlations between pairs of fMRI BOLD signals.

### 2.5. iEEG Data Analysis

For each participant, iEEG data were downsampled to 250 Hz. Artifact rejection involved three steps. First, outlier electrodes were identified based on the average log amplitude in one-minute segments across seven frequency bands computed using the demodulated band transform (DBT) (Kovach and Gander (2016)): delta (1-4 Hz), theta (4-8 Hz), alpha (8-14 Hz), beta (14-30 Hz), gamma (30-50 Hz), high gamma (70-110 Hz), and total power. Analytical amplitude measured in each band was z-scored across electrodes in each segment and averaged across segments. Electrodes with a mean z-score > 3.5 in any band were removed, including from further artifact rejection methods.

Second, intervals containing artifacts in the raw voltage traces were rejected on every electrode. We identified times when any electrode had extreme absolute raw voltage >10 SD for that electrode and marked as artifact the surrounding time until that electrode returned to zero voltage, plus an additional 100 ms before and after. Note that because we were measuring connectivity, any data interval identified on a single electrode was excluded for all electrodes. For each participant, if this procedure identified >1% of recording time as artifact, we optimized the total data kept for that participant (= electrodes × non-artifact time) by further excluding electrodes if retaining those electrodes caused more loss of data on the remaining electrodes via artifact rejection than they themselves contributed.

Third, we applied a specific additional noise criterion to eliminate brief power spikes in the high gamma band, a band that in some participants was particularly sensitive to noise in our recording environment. We excluded all intervals containing segments in which high gamma power averaged across electrodes exceeded the mean by >5 SD after excluding electrodes and times already removed in steps 1 and 2.

### 2.6. Multivariate Autoregressive Models

Multivariate autoregressive (MVAR) models of memory *p* describe the data in each of *M* channels as a weighted sum of *p* past values of data in all other channels and an error term. Let *y^m^*(*n*) be the value in channel *m, m* = 1,2,…, *M* at time *n, n* = 1, 2,…, *N* so we write

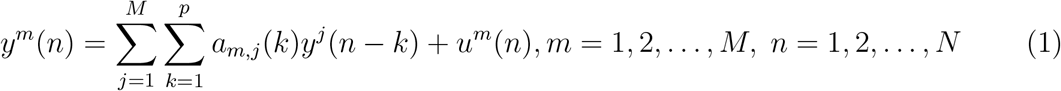

Here *a_m,j_*(*k*) is the weight applied to lag *k* of electrode *j* when modeling electrode m and *u^m^*(*n*) is the model error or innovations process, which is typically assumed to be temporally white and uncorrelated across electrodes. The number of data samples available is denoted by *N*.

The ordinary least squares problem for finding the model weights may be written

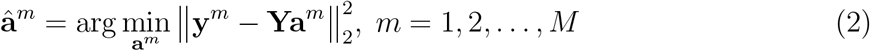

where

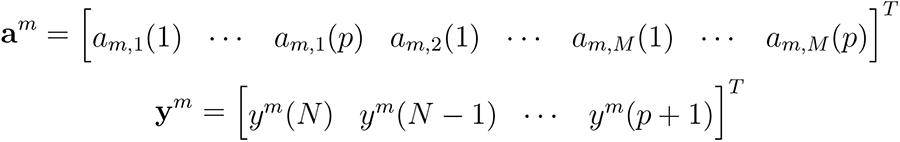

and

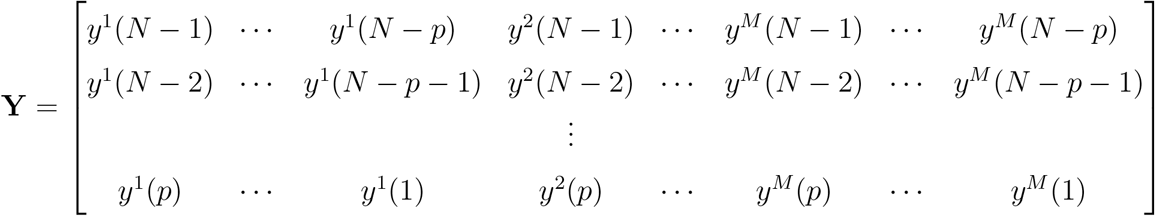

The solution to Eq. (2) is of the form

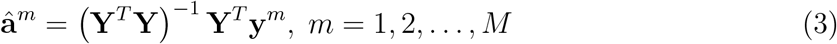

The prediction error variances are obtained from the MVAR weights as

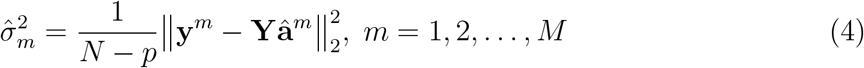

### 2.7. Ground Truth Networks and Data Simulation

The true underlying networks associated with the human iEEG data are not known. Hence, we use the data to estimate physiologically realistic ground truth MVAR models which are used to simulate data for assessing estimation performance as a function of data length. The ordinary least squares method produces poor quality models even with relatively long segments of human data, so we use ridge regression to regularize our ground truth models. That is, we solve

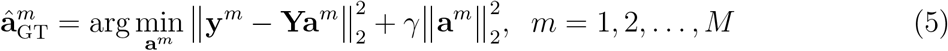

with

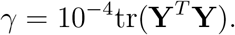

Define the prediction error signal

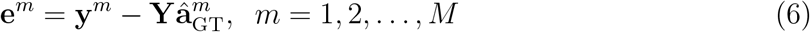

Thus, compute the error variance as

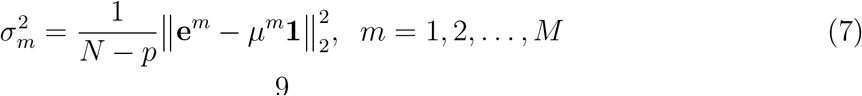

where the mean error in the channel is

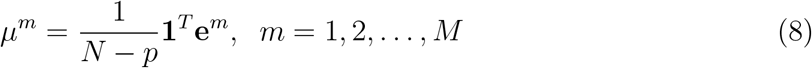

and **1** denotes an (*N* –*p*)-by-1 vector of ones. *μ^m^* is expected to be very small as preprocessing ensures the data is zero mean.

Nine resting state ground truth networks were fit in total, one network for each participant. Furthermore, in three subjects we fit ground truth networks to five sleep states (Wake, N1, N2, N3, REM), resulting in 15 additional ground truth networks. All 24 ground truth networks were estimated from 6 minutes of iEEG data. The simulated data for each ground truth network is obtained by computing Eq. (1) with an *u^m^*(*n*) a white noise process having variance 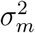 as defined in Eq. (7).

### 2.8. gLASSO and Weighted gLASSO Regularized Network Estimates

We used self-connected group LASSO regularization of Bolstad et al. (2011) as in Endemann et al. (2022). Define the weights connecting channel *j* to channel *m*

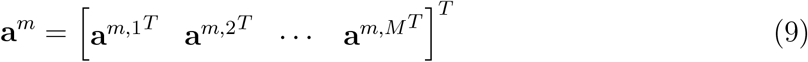

so that **a**^*m*^ is given by

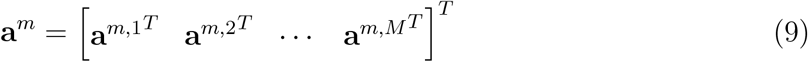

The gLASSO problem is then written as

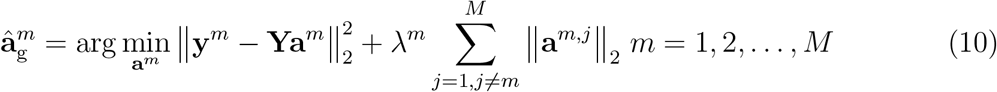

The gLASSO formulation in Eq. (10) penalizes all connections equally.

Prior information concerning the network model is incorporated by weighting the connection norms ||**a**^*m,j*^||_2_ inversely proportional to the expectation the connection is present in the model with a nonnegative scalar *w_m,j_*. This gives the weighted gLASSO (wgLASSO) problem

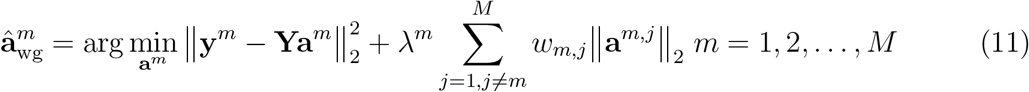

The MVAR model prediction error variances for either the gLASSO or wgLASSO models are obtained by substituting 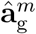 or 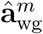, respectively, for 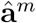 in Eq. (4). In the wgLASSO problem connections that have a high prior likelihood are assigned small or zero weights *w_m,j_* while connections that are unlikely are penalized more heavily with a relatively large weight.

In this work we used the fMRI Pearson correlation (PC) to determine the penalty weights *w_m,j_*. Intuitively, large values of PC between two regions indicate high likelihood of connection and thus those connections should be assigned small or zero weight in the penalty term. Let *c_m,j_* denote the Pearson correlation coefficient between electrode *m* and *j* where *c_m,j_* =*c_j,m_* as a non directed connectivity metric. The exact mapping between *c_m,j_* and the penalty weights is a design choice. We observed a log-linear relationship between the largest connection norms ||**a**^*m,j*^||_2_ of the ground truth networks and the corresponding |c_m,j_| in the resting state networks as illustrated in Figure 1. Motivated by this observation we adopted a conservative strategy whereby the connection norms are penalized based on ten raised to the power −*c_m,j_*|. This strategy is conservative as it avoids possible over penalization of the stronger connections - the upper decile shown in Figure 1B - at the expense of underpenalizing the weaker connections. Define

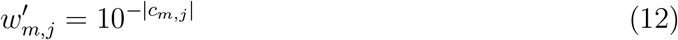

and then normalize 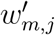 to the interval [0,1]

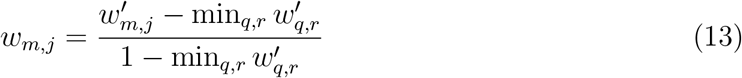

The normalization ensures that the connections with maximum PC in a particular participant are not penalized (*w_m,j_* = 0) while those with zero PC have a maximum penalty of unity.

Problems (10) and (11) are solved using the Expectation-Maximization algorithm of Bolstad et al. (2009). The hyperparameter λ^*m*^ is written in the form 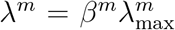 where 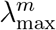 is the smallest value that results in the trivial solution of all zeros for the MVAR model weights. In particular,

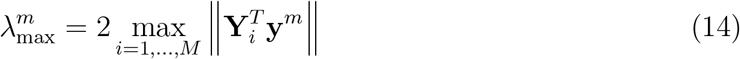

where **Y**_*i*_ is the (*N* – *p*) × *p* matrix block of **Y** that multiplies **a**^*m,i*^, for *i* = 1, 2,…, *M*. (See Eqs. (9) and (11).) The value *β^m^* is chosen using the following cross-validation procedure:

1. The entire training set is partitioned into five folds.
2. Select the *k*-th fold as the holdout set and train a model on the remaining folds for each candidate *β_i_* for every channel. All channels have the same candidate *β_i_* values to choose from, which consists of 10 logarithmically spaced values from 0.0001 to 1.
3. Use *k*-th fold holdout set to compute the mean square prediction error of each model.
4. Repeat steps two and three for all remaining *k* ∈{1, 2, 3, 4, 5}cases.
5. For each channel, set *β^m^* = *β_i_* for the *β_i_* value which corresponds to the lowest average mean square prediction error.

### 2.9. Performance Characterization

The MVAR model characterizes connection *m, j* in the network with the p parameters in the vector **a**^*m,j*^. Many different scalar parameterizations of connectivity strength have been proposed. In the body of this paper we use the broadband gPDC as described in Endemann et al. (2022), which is based on integrating the gPDC of Baccala et al. (2007) over frequency. The supplemental material demonstrates comparable results using the state-space Granger causality metric of Barnett and Seth (2015) to represent connectivity between electrodes.

We characterize the performance of each connectivity estimate via the cosine similarity between the recovered and ground truth connectivity matrices. The cosine similarity measures the angle, that is, the “alignment”, between two vectors **a** and **b**, and is defined to be

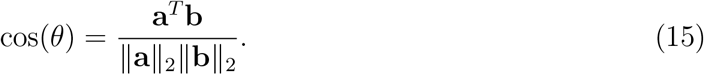

This similarity metric is applied to connectivity matrices **A** and **B** by stacking the columns of the matrices to form vectors and applying Eq. (15), which is equivalent to computing

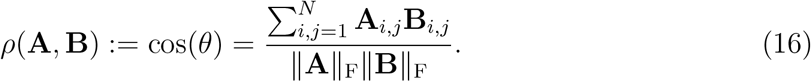

Note that the normalization in the denominator of both Eq. 15 and 16 means that the cosine similarity does not capture the effects of shrinkage associated with regularization.

We also use cross-validation to evaluate the normalized mean-square prediction error (NMSPE) on a subset of the data that was not used to estimate the MVAR model parameters. The NMSPE is computed for a given MVAR model 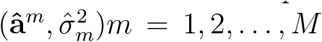 as

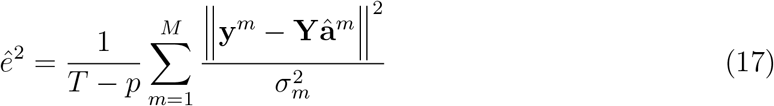

where **y**^*m*^ and **Y** have *T* – *p* rows and are constructed from the *T* samples of data held out from the model estimation process. Since we are simulating data, we choose *T* = 200, 000 to obtain a large data estimate of the NMSPE. Normalization by the ground-truth model error variance and use of a relatively large value for *T* ensures that *ê*^2^ has a minimum value of unity. The degree to which it exceeds unity reflects the excess prediction error associated with the estimated model.

## 3. Results

### 3.1. Comparison of estimation methods

We compared the performance of the gLASSO and wgLASSO estimators using MVAR model fits to data of varying lengths simulated from ground-truth models (see Methods). This is illustrated in Figure 2 for an exemplar participant with network size = 195 (Figure 2A), which shows the fMRI functional connectivity matrix that was used as the prior (Figure 2B), the matrix of penalty weights derived from this prior (Figure 2C), and the effective connectivity matrix of the ground truth model for this participant, measured as gPDC (Figure 2D). The corresponding SSGC ground truth connectivity is shown in Figure 3A. At each data length and for each estimator, performance was measured by calculating the cosine similarity (*ρ*) between the estimated and the ground truth connectivity matrices, and by measuring the MSE (*ê*^2^) between a holdout dataset and the model based on the estimated coefficients (note that the lowest possible MSE is 1). Performance of the wgLASSO estimator (Figure 2d) exceeded that of gLASSO (Figure 2e) for all but the longest data lengths investigated. For example, at a data length of *T* = 500 samples, the similarity of the recovered gPDC connectivity matrix to the ground truth for wgLASSO far exceeds that of gLASSO, and the MSE is much smaller. Similar results are obtained at *T* = 1000 samples (4 sec), but the performance difference is smaller. At *T* = 8000 samples (32 sec), performance is comparable for the two estimators. Comparable results were obtained for SSGC connectivity (Figure 3). Data for gPDC in all participants are shown in Supplementary Figure 4.

**Figure 2:**
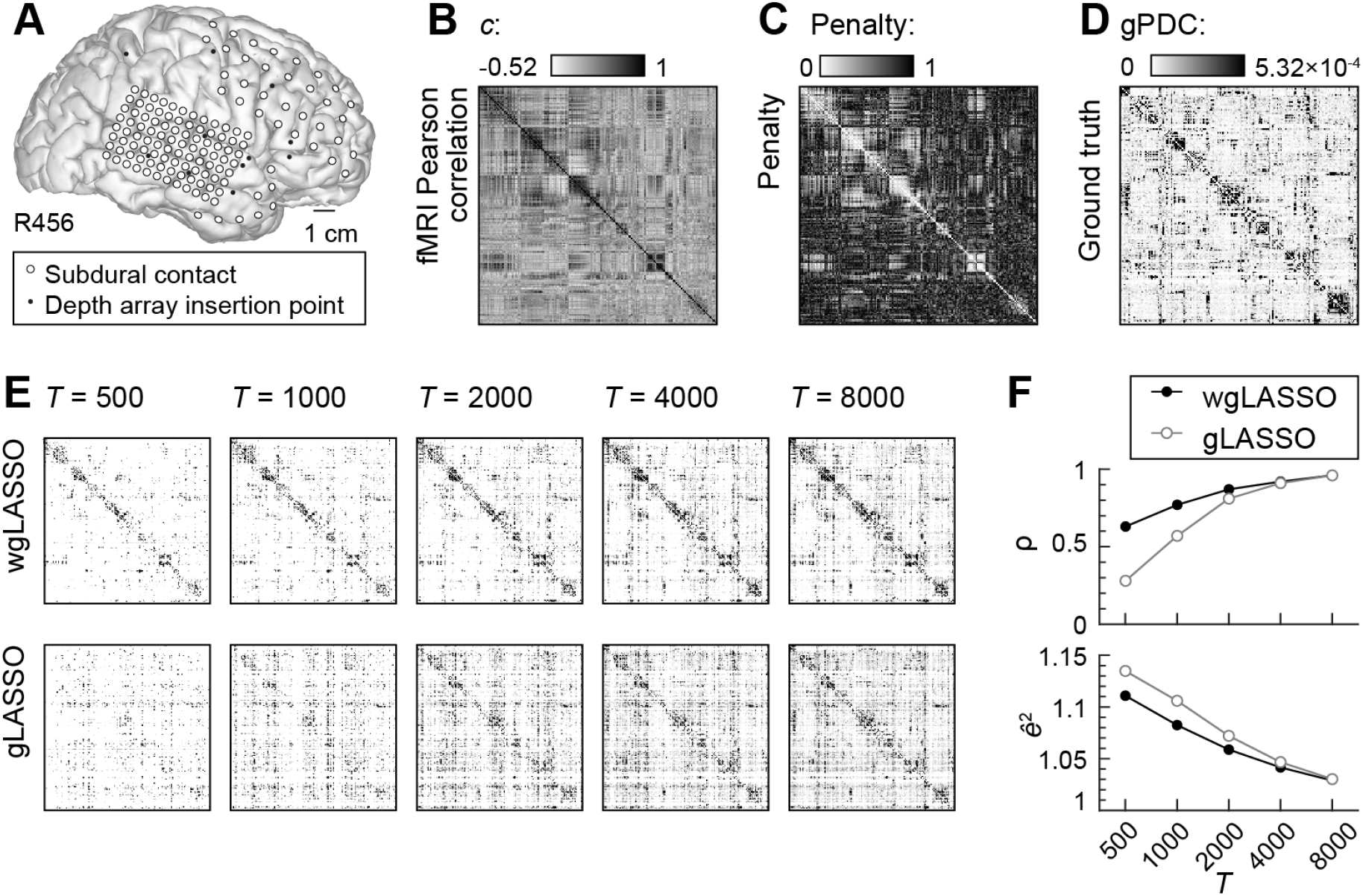
Comparison of model estimation using wgLASSO versus gLASSO for participant 456R. **(A)** Electrode coverage in subject 456R. **(B)** fMRI functional connectivity (Pearson correlation, *c*) matrix used as priors for wgLASSO. **(C)** Penalty matrix derived from the connectivity matrix in **A**. **(D)** gPDC connectivity matrix computed from ground truth model. **(E)** gPDC connectivity matrices from models estimated using wgLASSO (top *row*) or gLASSO (*bottom row*) fits to 5 different segment lengths ranging from *T* = 500 to 8000 samples. Each estimated connectivity matrix is averaged across five trials. **(F)** Estimator performance as a function of segment length for the data in **E**. *Top*: cosine similarity values (*ρ*) between estimated and ground truth gPDC matrices, averaged across the five trials; *bottom*: MSE (*ê*^2^) between the data and the model based on the estimated coefficients, also averaged across the five trials. For matrix plots in **B** - **D**, the darkest shading represents the 95th percentile of amplitude values.

**Figure 3:**
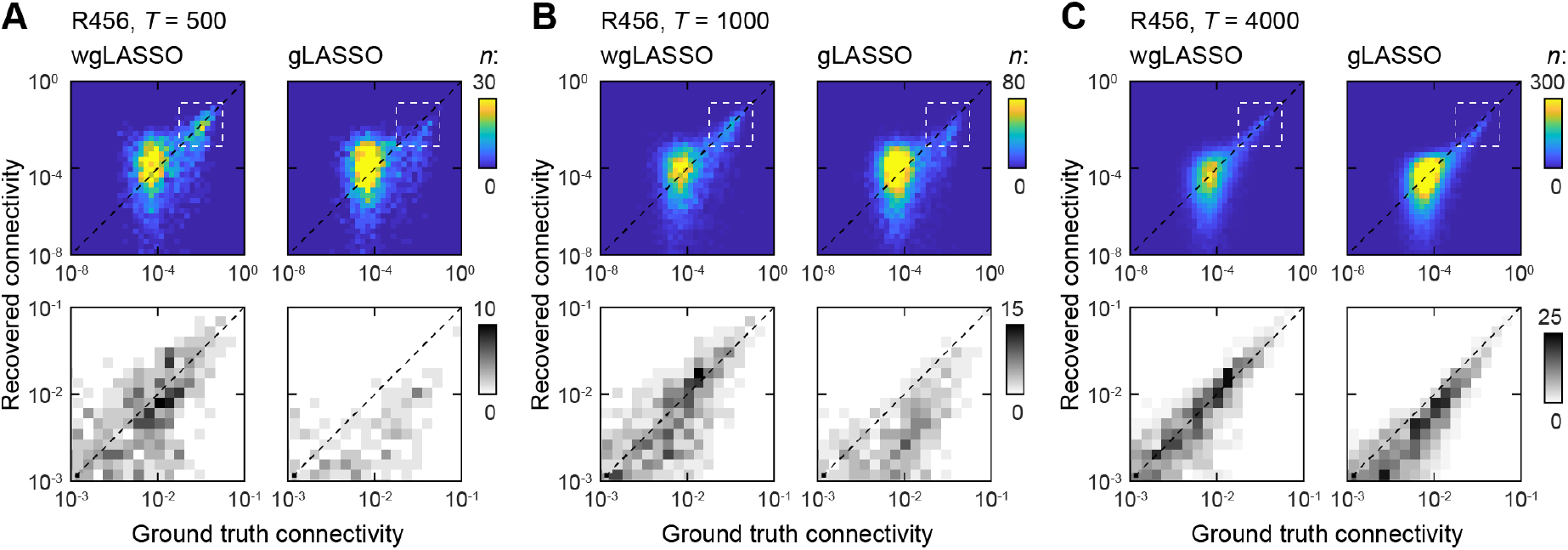
Comparison of recovered versus ground truth gPDC connectivity for participant 456R RS. Results for data lengths of **(A)** 500, **(B)** 1000, and **(C)** 4000 samples are shown; each data length consists of averaged gPDC across five trials.

Improved estimates of the strongest (and presumably most important) connections in the model using wgLASSO is illustrated most clearly in Figure 3, which shows 2-D histograms of recovered versus ground truth gPDC connectivity for three data lengths (same participant as Figure 2). For all three of the data lengths illustrated and for both estimators, the bulk of the recovered and ground truth gPDC values are small. However, more of the strongest connections are recovered using wgLASSO, and the estimates of these connections are more accurate, i.e., closer to the diagonal line. Note that the second row of each panel shows the histogram of the same data zoomed in to the strongest connections. The difference between wgLASSO and gLASSO in estimation accuracy for the strongest connections is diminished at T = 4000 samples, but the wgLASSO retains fewer of the weakest connections, as evidenced by the histograms of the full data range (Figure 3c, top row). Data for SSGC connectivity are shown in Supplementary Figure 5. Data from a second exemplar participant for gPDC are shown in Supplementary Figure 6.

Summaries of model performance across participants are shown in Figure 4. Here, we illustrate recovered gPDC connectivity measured as cosine similarity to the ground truth connectivity matrix (Figure 4A) as well as MSE (Figure 4B). Both assessments favor wgLASSO compared to gLASSO for all but the longest data length tested. Comparable results were obtained with SSGC connectivity (Supplementary Figure 7).

**Figure 4:**
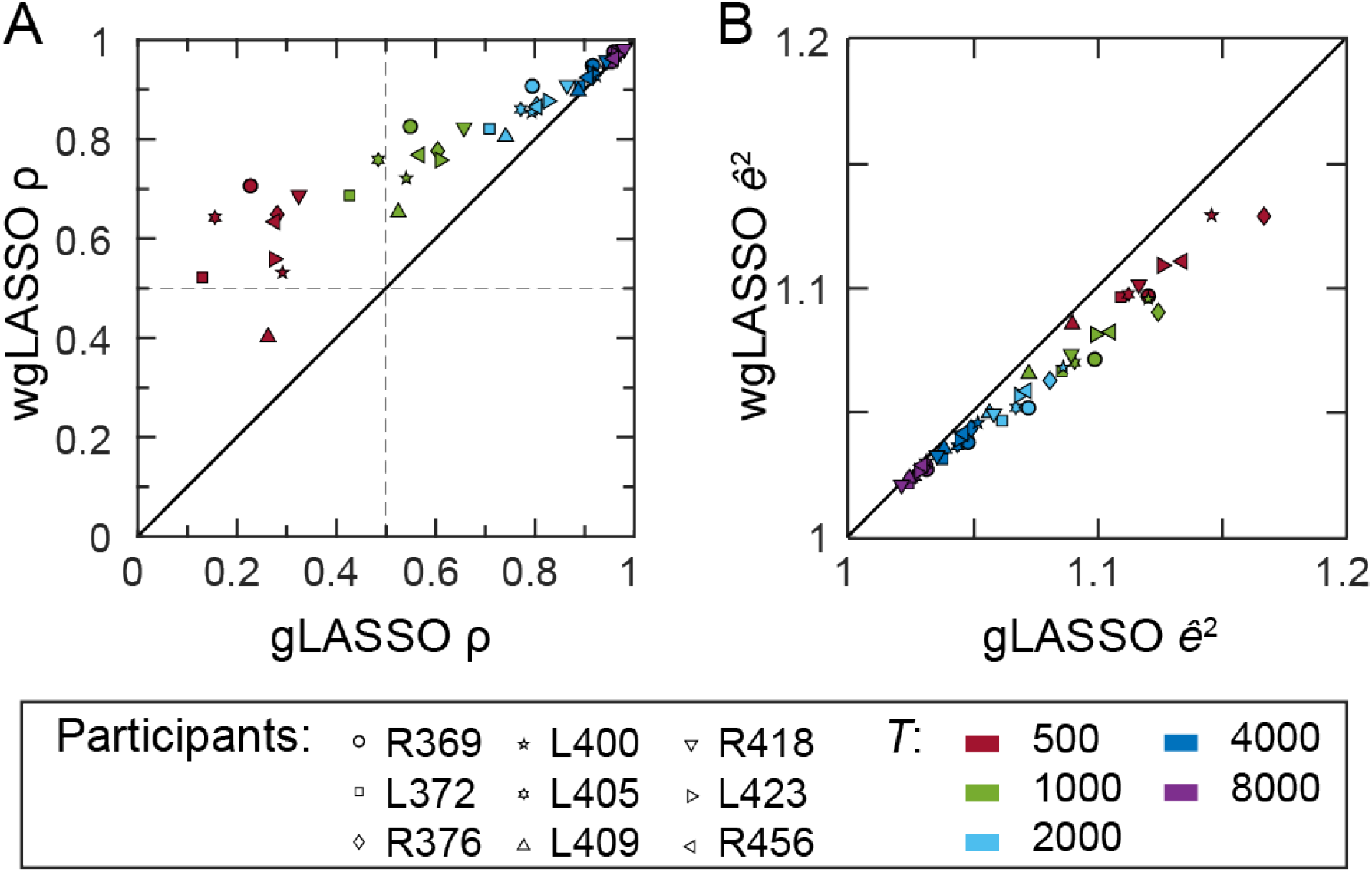
Summary of estimator performance across participants. **(A)** Cosine similarity computed between the estimated and ground truth gPDC connectivity matrices for wGLASSO plotted versus gLASSO estimators. **(B)** Similar to **A**, but for MSE. For both performance measures, the results for each participant and each data length are averaged across five trials.

As indicated by the 2-D histograms in Figure 3 and Supplementary Figures 5 and 6, although gLASSO and wgLASSO have increasingly similar performance as the data length increases, the wgLASSO estimator uses a more parsimonious model. This is illustrated in Figure 5, which shows the percent of connections pruned for wgLASSO compared to gLASSO. Especially at the longest data lengths, wgLASSO estimates were consistently sparser compared to gLASSO.

**Figure 5:**
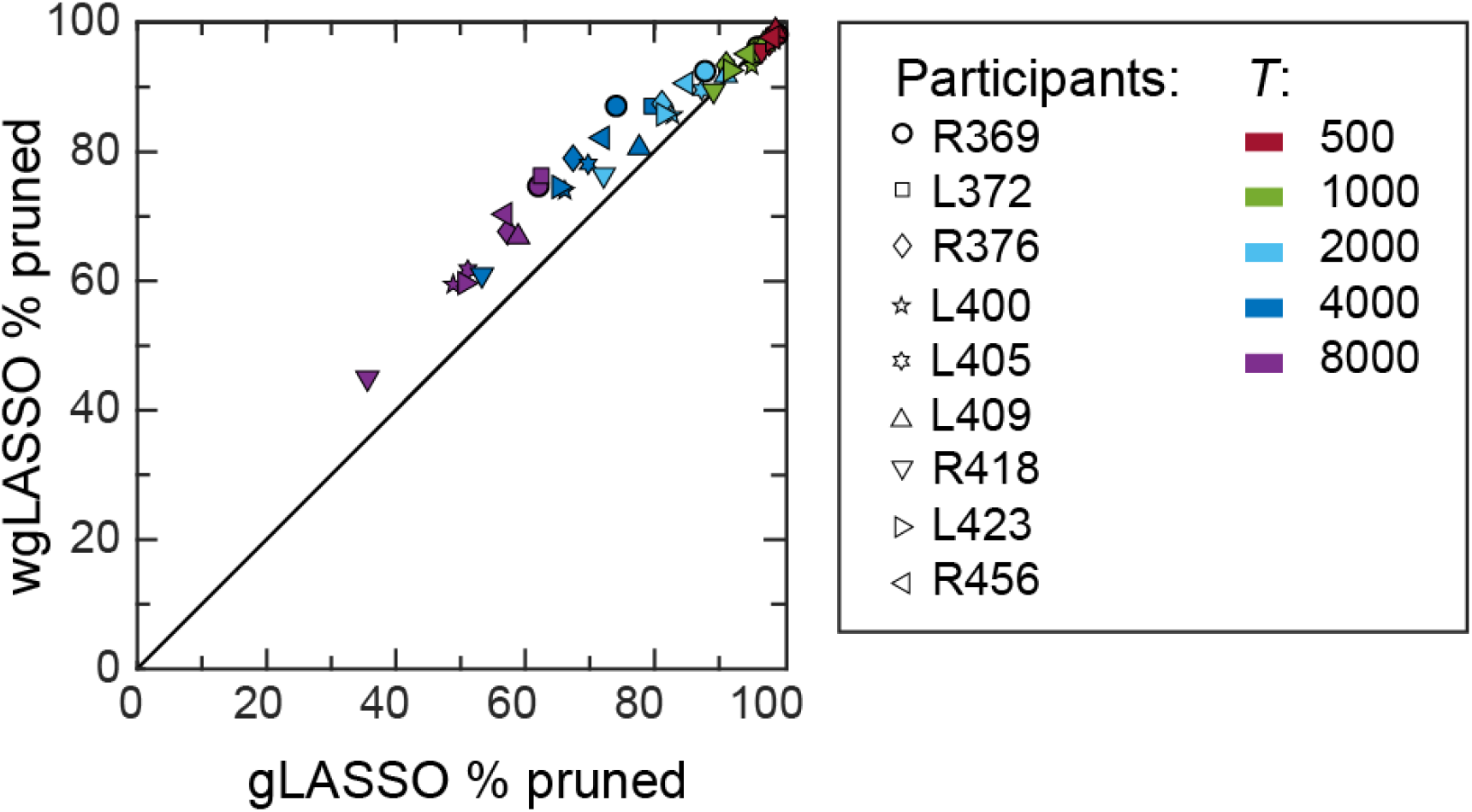
Summary assessment of the the percentage of pruned connections in the gPDC recovery matrices using gLASSO and wgLASSO methods for all nine RS participants. The (*i,j*) – *th* connection is considered pruned if the (*i,j*) – *th* element of a method’s gPDC recovery matrix is zero and the (*i,j*) – *th* element of the ground truth gPDC matrix is non-zero. The results for each participant are averaged across five trials.

### 3.2. Robustness of estimators across brain and behavioral states

The priors for wgLASSO are derived from resting state fMRI data collected during the day when participants were awake. We have shown that wgLASSO improves model estimates when applied to iEEG data collected under similar conditions, i.e., daytime wake. It is often the case, however, that iEEG data is collected under very different conditions than the fMRI priors, and the question arises as to whether these priors would be useful when applied to such data. We addressed this question by applying wgLASSO to data recorded during sleep and staged using polysomnography (Banks et al. (2020)). We chose 3 of the 9 participants included in the resting state dataset presented above (376R, 418R, 423L). We used the same fMRI priors as for the resting state dataset. We evaluated the performance of the wgLASSO estimator when applied to data simulated from ground truth models that were derived from data recorded during five identified sleep stages (wake, REM, N1, N2, N3).

Figure 6 depicts 2-D histograms of estimated versus ground truth connectivity for both gLASSO and wgLASSO in wake, N3, and REM based on *T* = 1000 samples. Figure 8 depicts the fMRI connectivity (A), penalty weights (B), and compares ground truth, wgLASSO, and gLASSO connectivity estimates for the same three states. The fidelity of estimates obtained with gLASSO varied across sleep stages, likely because *T*=1000 samples is on the threshold of failure for gLASSO and the signal to noise ratio varies across states. As with the resting state data, wgLASSO yielded improved recovery of the strongest connections while producing sparser models, and these results were more consistent across sleep stages compared to gLASSO. For example, compare Figure 6B showing data from N3 sleep to 6A and 6C showing data from wake and REM sleep, respectively.

**Figure 6:**
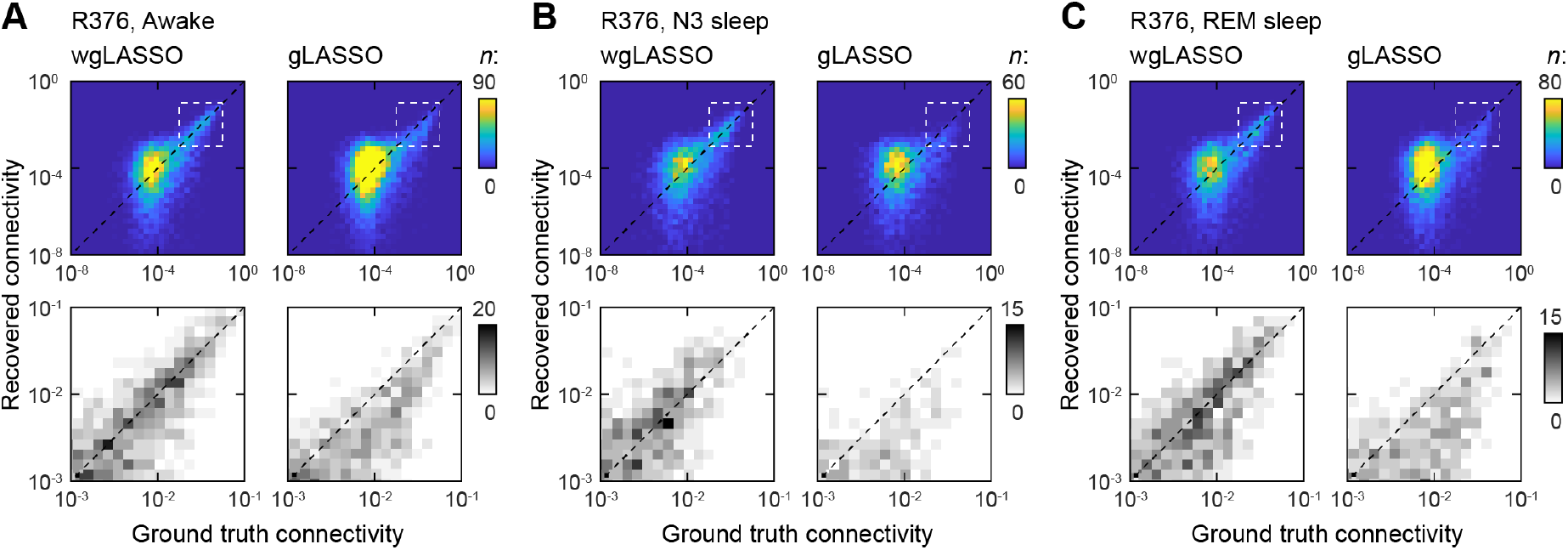
Comparison of recovered gPDC connectivity against ground truth gPDC connectivity for data recorded during sleep for participant 376R. Same data as in Figure 8. Three of the five sleep states are shown. **(A)**. Awake. **(B)**. N3. **(C)**. REM. Each estimated gPDC matrix is averaged across five trials for each method; 1000 training samples were used to fit the estimated models.

**Figure 7:**
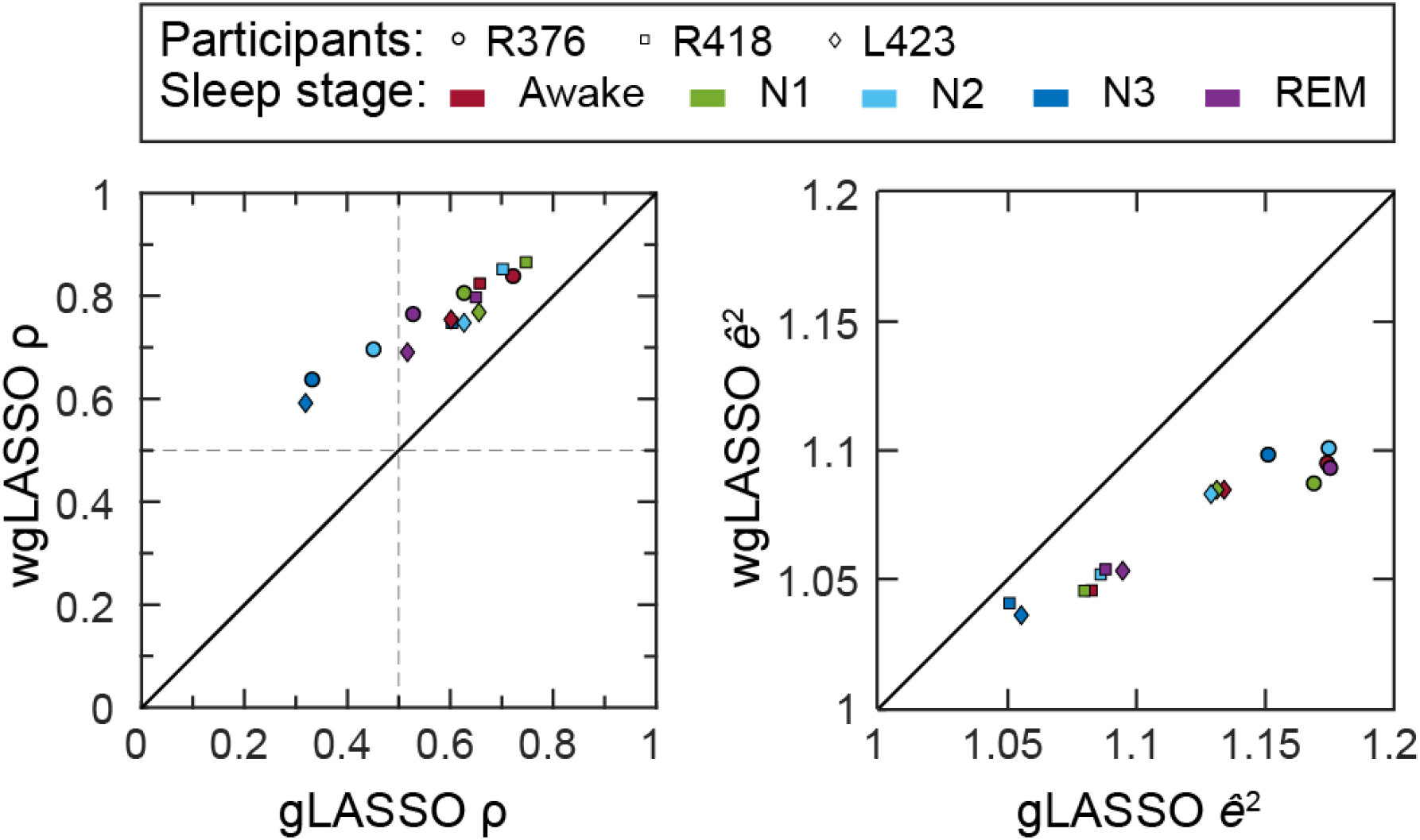
Summary assessment of gPDC recovery (left) and model performance (right) using the gLASSO and proposed wgLASSO methods for all three sleep participants. Cosine similarity was computed between the estimated gPDC and ground truth gPDC connectivity matrices. One-step MSE on a holdout set was used to determine model performance for each method, and the ratio is computed between them. The results for each participant are averaged across five trials assuming *T* = 1000 samples.

**Figure 8:**
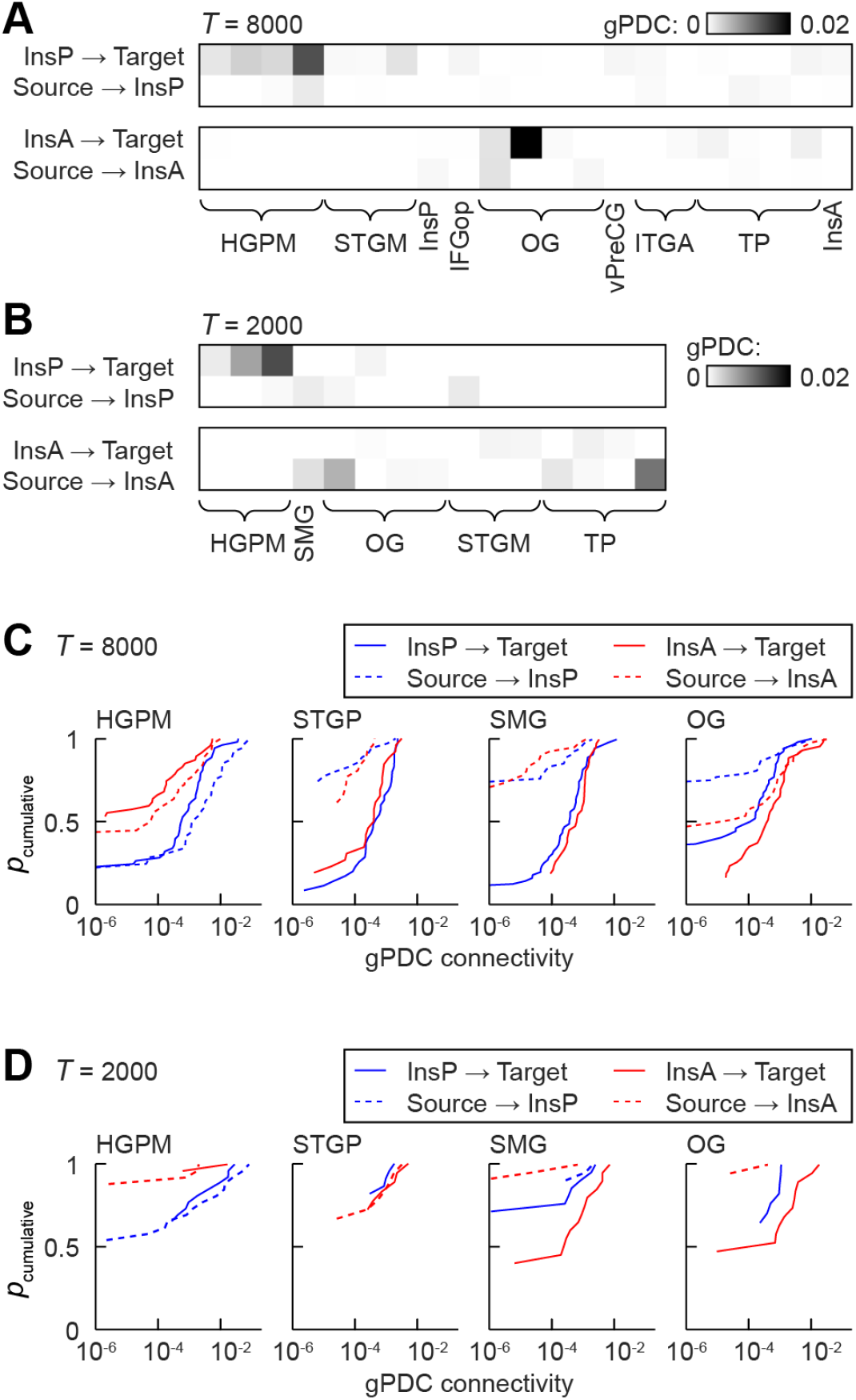
Connectivity of the anterior and posterior insula. **(A)**. Matrix showing gPDC connectivity between single (and only) recording sites in InsP and InsA, and recording sites in 9 regions of interest in 423L. The selected regions of interest were those with electrode coverage in all 7 participants and that had non-zero gPDC values in this participant. *T* = 8000 samples. **(B)**. As in A, for *T* = 2000 samples. Note that because of the shorter data length, the estimated model is sparser and fewer connections had non-zero gPDC values. **(C)**. Cumulative distribution functions of gPDC connectivity across recording sites and participants for InsP (blue lines) and InsA (red lines), for projections to the indicated target region (solid lines) and from the indicated source region (dashed lines). *T* = 8000 samples. **(D)**. As in C, for *T* = 2000 samples. (Abbreviations. HGPM: postero-medial portion of Heschl’s gyrus; STGM: medial portion of the superior temporal gyrus; InsP: insular cortex, posterior portion; IFGop: inferior frontal gyrus, pars opercularis; OG: orbitofrontal gyrus; vPreCG: precentral gyrus, ventral portion; ITGA: inferior temporal gyrus, anterior portion; TP: temporal pole; InsA: insular cortex, anterior portion)

The summary of results across all three participants are shown in Figure 7. For all participants, recovery of ground truth connectivity and model performance are superior for wgLASSO compared to gLASSO. Improvement in recovery and performance was greater for stages showing poorest results with gLASSO. The fMRI prior resulted in improved recovery even though obtained from a different behavioral state.

### 3.3. Connectivity estimates from model fits to recorded data

The results presented so far are based on model fits to simulated data, which enabled comparison to a ground truth condition. A real-world test of model performance involves model fits to recorded data, which are not guaranteed to conform to the assumptions of MVAR models, likely contain nonlinearities, and may contain noise that is not eliminated in preprocessing. In this case, there is no ground truth to serve as comparator, but as in Endemann et al. (2022) we compared connectivity for two brain regions, posterior and anterior subdivisions of the insula (InsP, InsA). These neocortical regions are anatomically adjacent and strongly interconnected, but have divergent connectivity profiles with the rest of the brain and are functionally distinct (Augustine (1996) Cauda et al. (2011), Cloutman et al. (2012), Zhang et al. (2019), Endemann et al. (2022)). InsP is a sensory area, receiving both interoceptive and exteroceptive input from the periphery and other cortical regions. By contrast, InsA is a higher order cortical region, with strong connectivity to prefrontal cortex and limbic structures. Our expectation, then, is that InsP will be more strongly connected with primary sensory regions such as core auditory cortex (posteromedial Heschl’s gyrus, HGPM), and InsA will be more strongly connected with higher order areas such as the prefrontal cortical region orbitofrontal gyrus (OG). Connectivity estimated from MVAR model fits aligned with this expectation.

Seven of the nine participants in the RS dataset had electrodes in both InsP and InsA. Using wgLASSO, models were fit in these seven participants to data from all recording sites (see Table 1) using two different data lengths, 2000 and 8000 samples, and connectivity (gPDC) calculated. Results for one participant are shown in Figure 8 A,B (*T* = 8000 and *T* = 2000 samples, respectively). The vertical axis in each connectivity matrix shows the four classes of connections, i.e. gPDC from and to InsP and InsA. The subset of source and target regions with electrode coverage in all 7 participants are indicated on the horizontal axis, and are ordered roughly hierarchically as in our previous work, with HGPM on the left and higher order cortical regions on the right. Heavier connectivity between InsP and lower order regions is illustrated by the denser grey scale values on the left side of the connectivity matrix. Heavier connectivity between InsA and higher order regions is illustrated by the denser grey scale values on the right side of the connectivity matrix. Note that although the model fits for *T* = 2000 samples are sparser, the same pattern is recapitulated.

Connectivity data are summarized across participants for four representative regions of interest in Fig. 8 CD (*T* = 8000 and *T* = 2000 samples, respectively), which shows cumulative distribution functions (cdfs) of gPDC values for InsP (blue) and InsA (red) across recording sites and participants. As in the single participant example above, regions of interest are ordered roughly hierarchically, with lower order areas on the left. Stronger connectivity is indicated by a rightward shift of the cdf curve along the horizontal axis in each panel. For HGPM, the cdfs for InsP are shifted to the right relative to InsA, indicating stronger connectivity with core auditory cortex. By contrast, for OG the opposite pattern is apparent. Intermediate regions such as superior temporal gyrus, posterior portion (STGP) and supramarginal gyrus (SMG) show intermediate patterns. Again, even though the model fits for T = 2000 samples are sparser, the same pattern is recapitulated. Overall, these results strongly suggest that the wgLASSO approach reliably captures essential effective connectivity information in the short data length regime.

## 4. Discussion

Previously, we demonstrated that gLASSO is an effective method relative to ordinary least squares for significantly reducing the data required to estimate MVAR models of brain networks with large numbers of recording sites (Endemann et al. (2022)). The gLASSO encourages sparse connectivity by penalizing the two-norms of the weight vectors associated with all connections. Hence, the gLASSO treats the connections between all brain regions equally. The wgLASSO offers a straightforward mechanism for differential penalization of connections based on prior expectations, such as provided by participant-specific structural or functional connectivity measures. Connections that are weak in these prior measures are less likely to be significant and may be assigned a relatively large penalty weight, while those that are likely may be assigned a small or zero penalty weight. Importantly, incorporation of prior information via the wgLASSO further reduces the data required to estimate MVAR models for large numbers of recording sites. Our simulation studies using neurophysiologically realistic ground-truth models show that the fMRI-based wgLASSO method proposed in this work reduces the data requirements by an additional factor of approximately two relative to that of the gLASSO. The results in Figs. 2, 3, 4 and 7 imply our fMRI-based wgLASSO approach is able to reconstruct important connectivity features in networks of 200 channels with 4-8 seconds of data.

The degree to which prior information can reduce data requirements depends on the nature and quality of the prior information and the manner in which it is incorporated into the estimation algorithm. fMRI and iEEG measure neural activity on different time scales, and although functional connectivity measures derived from the two modalities are related, the correlations are modest (Hacker et al. (2017)). In the current study, the specific circumstances of the recording sessions for fMRI and iEEG data were distinct: fMRI data were collected before the electrode implantation surgery, while the iEEG data were recorded post-operatively. Furthermore, the PC values used as priors are not directed, while the iEEG MVAR model represents directed connections. Indeed, as shown in Figure 1A, the relationship between the ground-truth model connection norms and PC values is not specific - there is a wide range of MVAR model connection norms associated with each PC value. Hence, incorporation of prior fMRI information must be done in a flexible manner to accommodate differences between iEEG and fMRI connectivity.

The fMRI-based wgLASSO approach proposed here readily accommodates the differences between fMRI and iEEG via differential penalization of connection norms. First, the prior information is not applied at the level of the MVAR model coefficient values, mitigating the differences in time scale between fMRI and iEEG measures. Further, the wgLASSO in Eq. (11) is a soft approach in that it trades the reduction in squared error associated with a particular connection against the penalty associated with that connection. Connections that are strongly penalized can still be present in the MVAR model if they contribute to a proportionally large reduction in squared error. On the other hand, if two connections contribute equally to the squared error, the connection that is least penalized will be favored. Finally, the wgLASSO approach presented here is flexible, and can easily accommodate other types of prior information, as as from diffusion tensor imaging (Zhu et al. (2013), Belaoucha and Papadopoulo (2020)) or electrical stimulation tract tracing (Rocchi et al. (2021)).

The conservative strategy proposed here for choosing the penalty weights applied to each connection is motivated by the nonspecific relationship between the MVAR model connection norms and PC values illustrated in Fig. 1A. A logarithmic relationship between PC values and the largest connection norms in the ground truth models was observed across all participants (Fig. 1C, D) so we chose the penalty for all connections associated with a particular PC value c based on 10^-|*c*|^ normalized to the interval [0, 1]. Connections with PC values of 1 are not penalized, while those with PC value zero receive maximum penalty of 1. Hence, connections with similar PC values are penalized comparably and the wgLASSO cost function will favor connections that give the largest squared error reduction for a given connection norm. Deriving the penalty based on the relationship between PC and connections with the largest norms minimizes the risk of over penalization. Note that more aggressive penalization strategies are possible and would likely further reduce data requirements at the expense of forcing the estimate to match the prior. That is, over penalization could lead to important connections being discouraged by the wgLASSO.

The relative benefits of the proposed wgLASSO approach over gLASSO is evident in all participants for multiple performance metrics. The cosine similarity metric for the wgLASSO gPDC (Fig. 4A) and state-space Granger causality measure (Supplementary Fig. 7) is consistently higher than that for the gLASSO estimates except at the very longest data lengths considered (8000 samples or 32 seconds). The wgLASSO models have lower one-step prediction errors than the gLASSO models (Fig. 4A) over the same range of data lengths. Even at these long data lengths, however, we note that the wgLASSO models are sparser, i.e., use fewer connections, than the gLASSO models (see Fig. 5) while achieving comparable model performance. Thus, the wgLASSO method results in a more parsimonious model for the measured data.

Furthermore, we show that the improved model performance holds even for data recorded under very different conditions than the fMRI priors. We already noted the time separation with fMRI recorded pre-operatively and iEEG recorded post-operatively. We also evaluated model performance when iEEG data were recorded during different stages of overnight sleep, when cortical networks can exhibit substantially different connectivity profiles compared to the daytime wake conditions of the fMRI scans (Banks et al. (2020)). Even so, the proposed penalization strategy using wgLASSO still outperforms gLASSO across these distinct behavioral states as shown in Figures 6 and 7, and Supplementary Figure 8. This robustness is likely due in part to the soft way the wgLASSO incorporates prior information and conservative manner in which we chose the penalty weights.

The gLASSO consistently shows evidence of shrinkage in the largest coefficients, i.e., the estimated connection strength is less than the ground truth value (see Fig. 3 and Supplementary Figs. 5 and 6), particularly with relatively short data lengths. This shrinkage is a well-known effect associated with penalized least-squares formulations and is sometimes addressed by a debiasing step as noted in Endemann et al. (2022). Shrinkage is much less evident in the largest coefficients with the wgLASSO method because these larger connectivity values are associated with the largest PC values, which are mapped to penalty weights at or near zero. Hence, the wgLASSO approach proposed here tends to be less affected by shrinkage and subsequent debiasing steps are likely unnecessary.

We note the limitations of the ground truth models employed here. Ground truth models are necessary to quantitatively evaluate estimation methods and compare performance. However, the strength of our conclusions is limited by degree to which our ground truth models, which are linear and stationary, create representative data. We note that the fMRI prior employed here is independent of the ground truth models, and its success provides independent evidence that these models are reasonable. The wgLASSO approach also yielded results consistent with known brain connectivity profiles when applied to recorded data rather than data simulated from GT models (Figure 8). In the example shown comparing connectivity to posterior and anterior insula, lower order sensory areas had stronger connectivity to posterior insula, consistent with its role in sensory processsing. By contrast, higher order areas had stronger connectivity to anterior insula, consistent with its role as a gateway to limbic cortex (Augustine (1996) Cauda et al. (2011), Cloutman et al. (2012), Zhang et al. (2019), Endemann et al. (2022)).

The wgLASSO approach is motivated by concerns over stationarity, which diminish with decreasing data length. Over sufficiently short time scales a stationary approximation to brain data is expected to be effective at providing insight into how brain network organization evolves over time. In addition to enabling investigation of brain dynamics per se, the approach presented here can be leveraged for analysing connectivity changes during an experiment with distinct epochs, such as a behavioral task. For example, 0.1 sec time windows could be concatenated over many trials to investigate connectivity changes that evolve over a delay period during a working memory task. Thus, the wgLASSO approach has the potential to inform investigations of brain network reorganization underlying rapid behavioral adaptations.

## Supplementary Figures

**Supplementary Figure 1:**
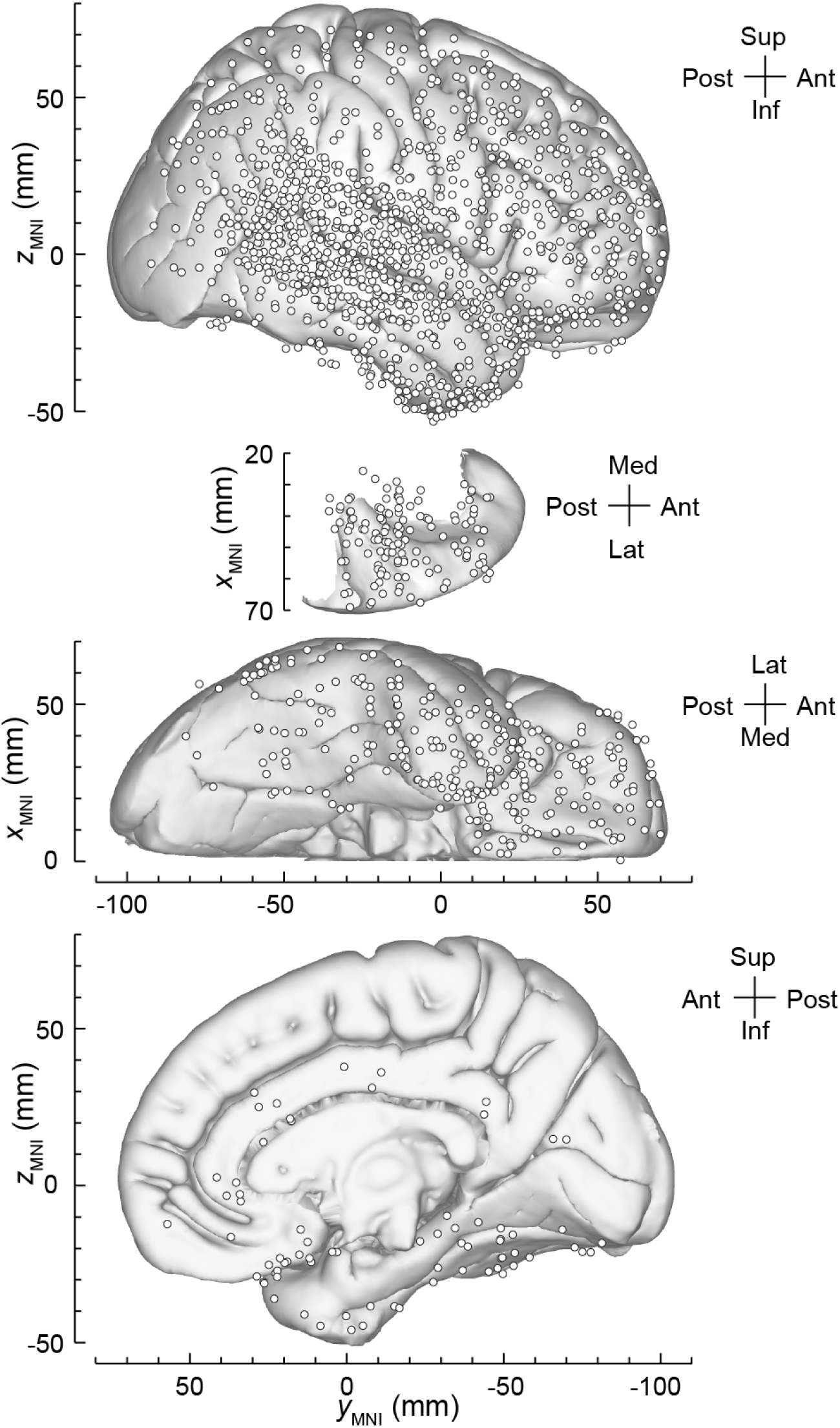
Summary of electrode coverage in 9 participants recorded during daytime waking periods. Locations of recording sites are plotted in Montreal Neurological Institute (MNI) coordinate space and projected onto the Freesurfer average template brain for spatial reference. Projections are shown on the lateral, top-down (superior temporal plane), ventral and mesial views (top to bottom). Recording sites over orbital, transverse frontopolar, inferior temporal gyrus and temporal pole are shown in both the lateral and the ventral view. Some recording locations in regions inaccessible to these views (e.g., hippocampus, amygdala) are not shown. Recording locations in participants with electrodes implanted into the left hemisphere have been reflected onto the right hemisphere for visualization purposes.

**Supplementary Figure 2:**
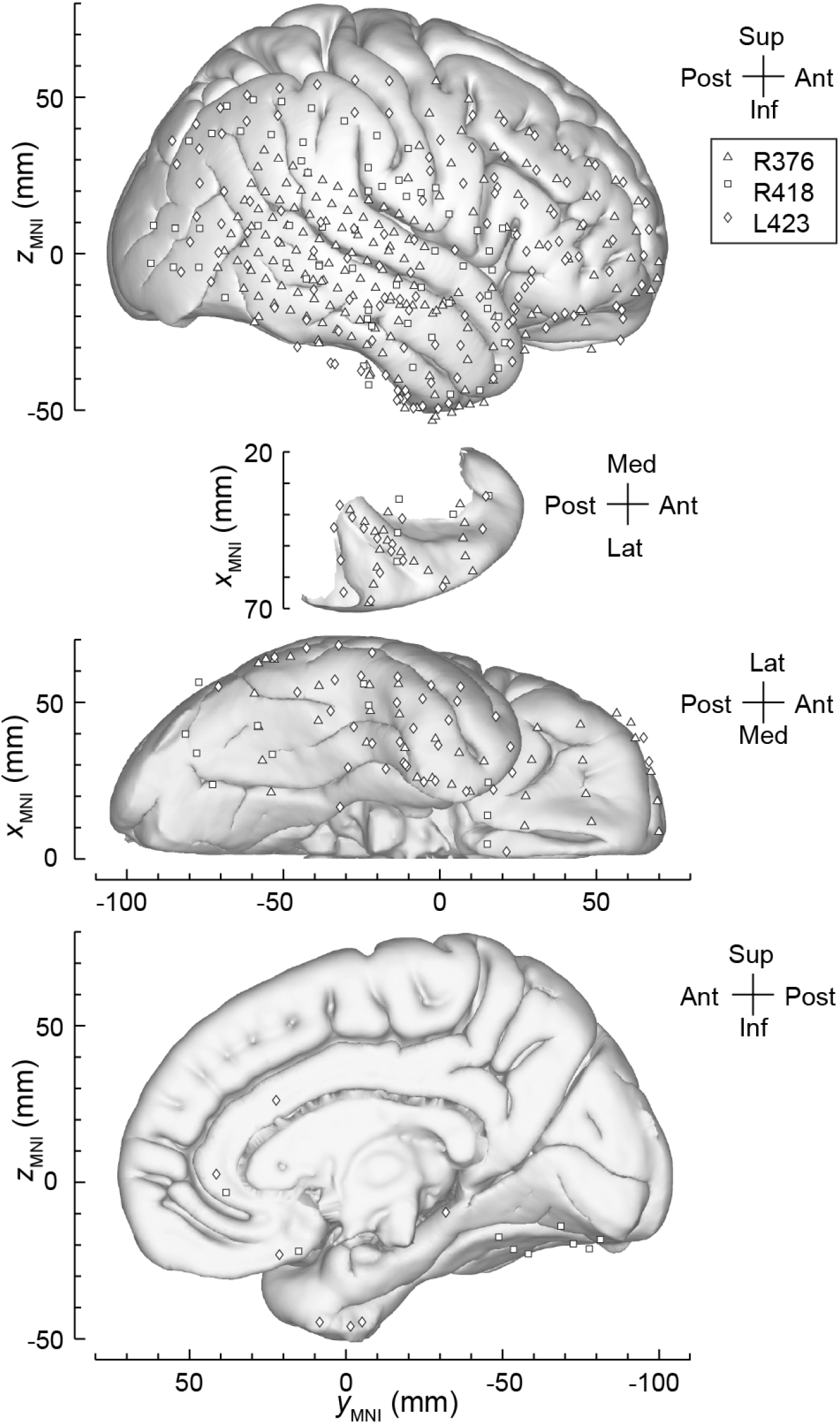
Summary of electrode coverage in 3 participants recorded during overnight sleep. See Supplementary Figure 1 for details.

**Supplementary Figure 3:**
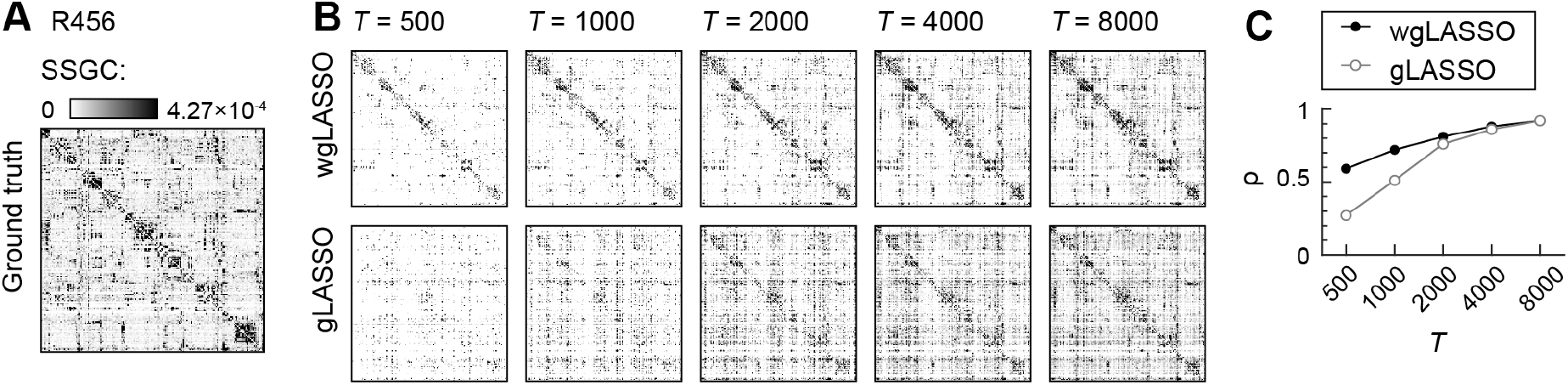
Comparison based on SSGC of model estimation using wgLASSO versus gLASSO for participant 456R. **(A)** SSGC connectivity matrix computed from ground truth model fit using ridge regression to 6 minutes of resting state data. **(B)** SSGC connectivity matrices from models estimated using wgLASSO (*top row*) or gLASSO (*bottom row*) fits to 5 different segment lengths ranging from *T* = 500 to 8000 samples. Each estimated connectivity matrix is averaged across five trials. **(C)** Cosine similarity values between estimated and ground truth SSGC matrices shown in **B**, averaged across the five trials (*ρ*), plotted versus segment length. See Figure 2**B** & **C** for fMRI functional connectivity and penalty matrices used for wgLASSO.

**Supplementary Figure 4:**
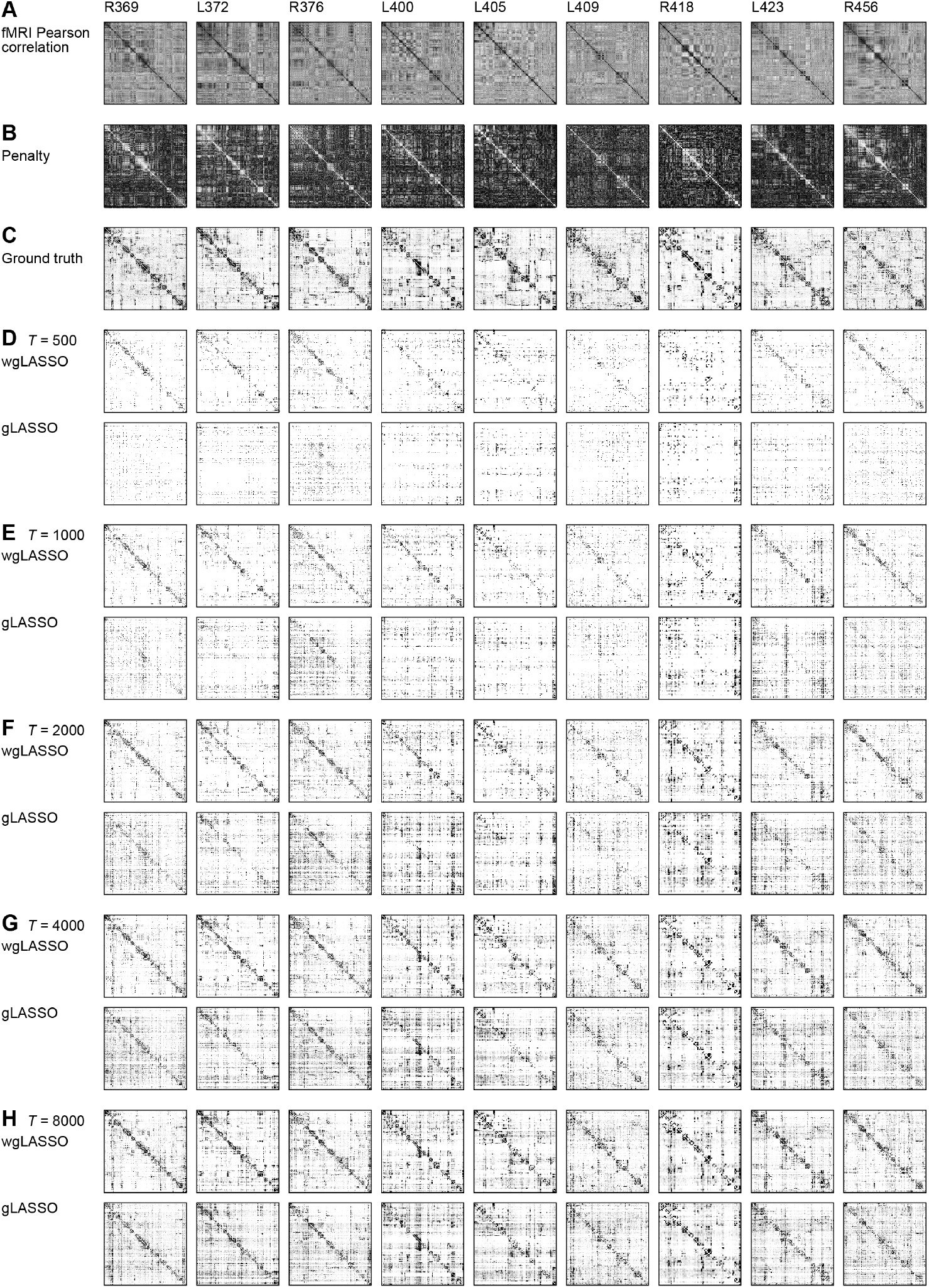
Recovery of gPDC connectivity using gLASSO and wgLASSO methods for all RS participants. Each column is a different participant. **(A)**fMRI Pearson correlation matrices. **(B)** Penalty matrices used for wgLASSO. **(C)** gPDC connectivity matrices of ground truth models. **(D)** - **(H)** gPDC matrices from models estimated using wgLASSO (*top row*) or gLASSO (*bottom row*) fits to segment lengths ranging from *T* = 500 to 8000 samples. Each estimated connectivity matrix is averaged across five trials.

**Supplementary Figure 5:**
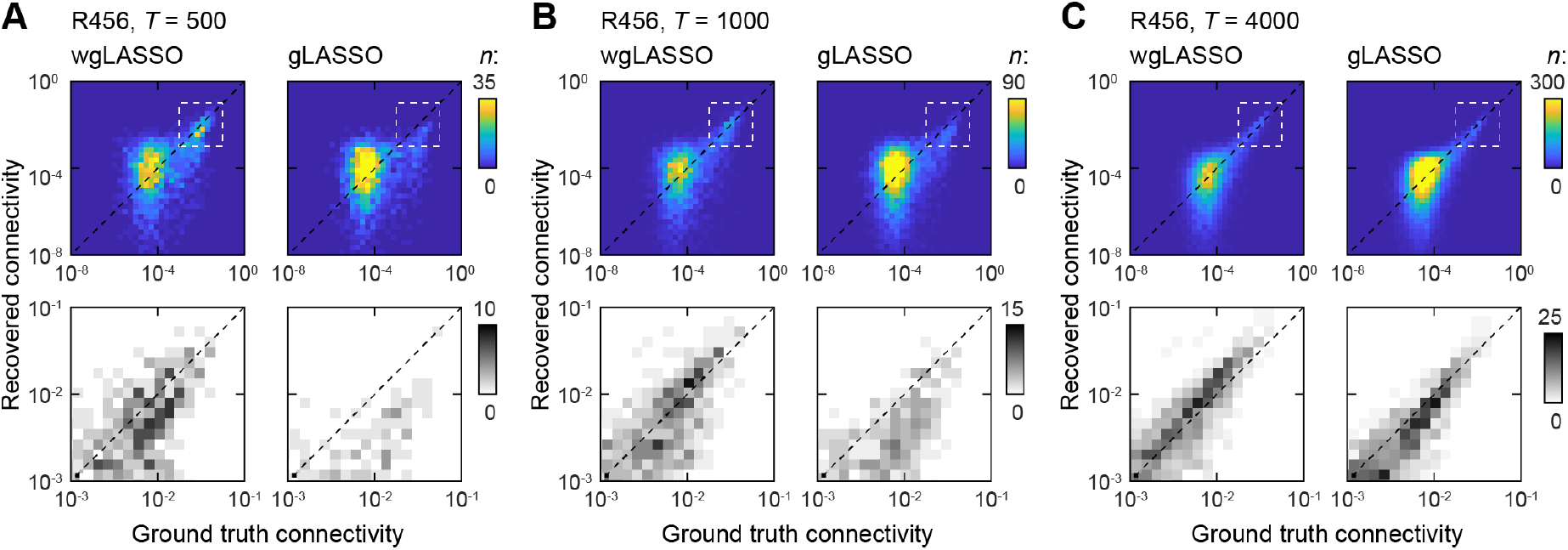
Comparison of recovered versus ground truth SSGC connectivity for participant 456R RS. Results for data lengths of **(A)** 500, **(B)** 1000, and **(C)** 4000 samples are shown; each data length consists of averaged SSGC across five trials.

**Supplementary Figure 6:**
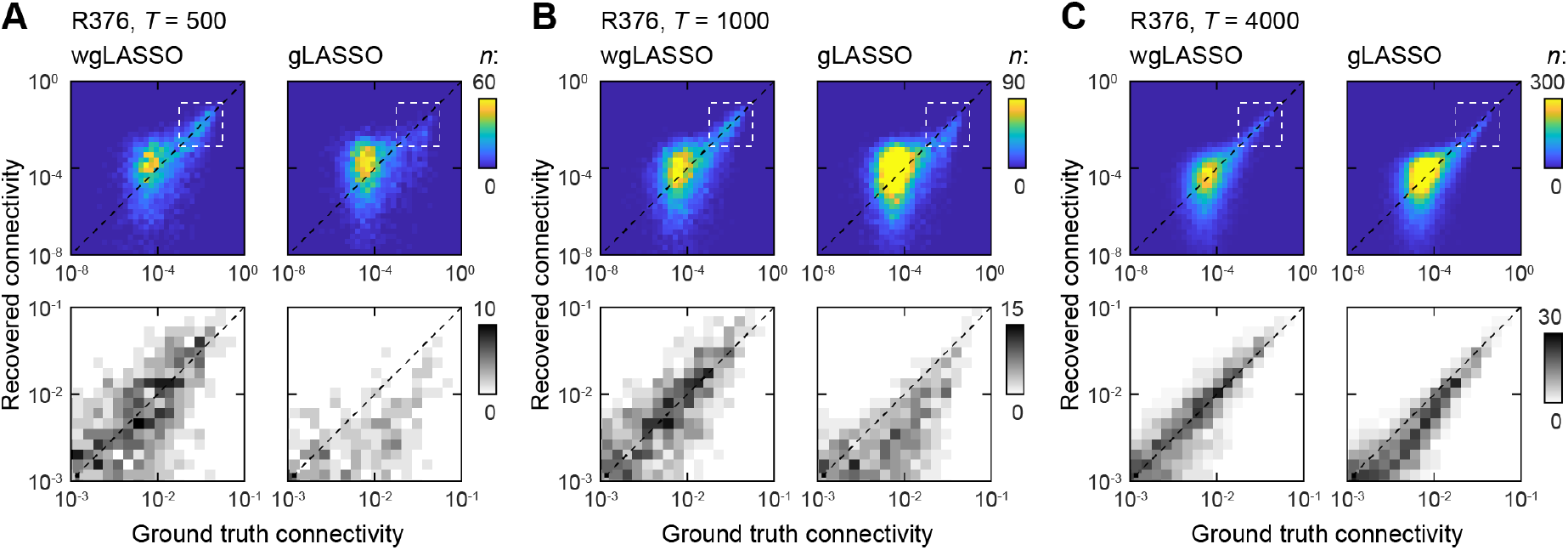
Comparison of recovered versus ground truth gPDC connectivity for participant 376R RS. Results for data lengths of **(A)** 500, **(B)** 1000, and **(C)** 4000 samples are shown; each data length consists of averaged gPDC across five trials.

**Supplementary Figure 7:**
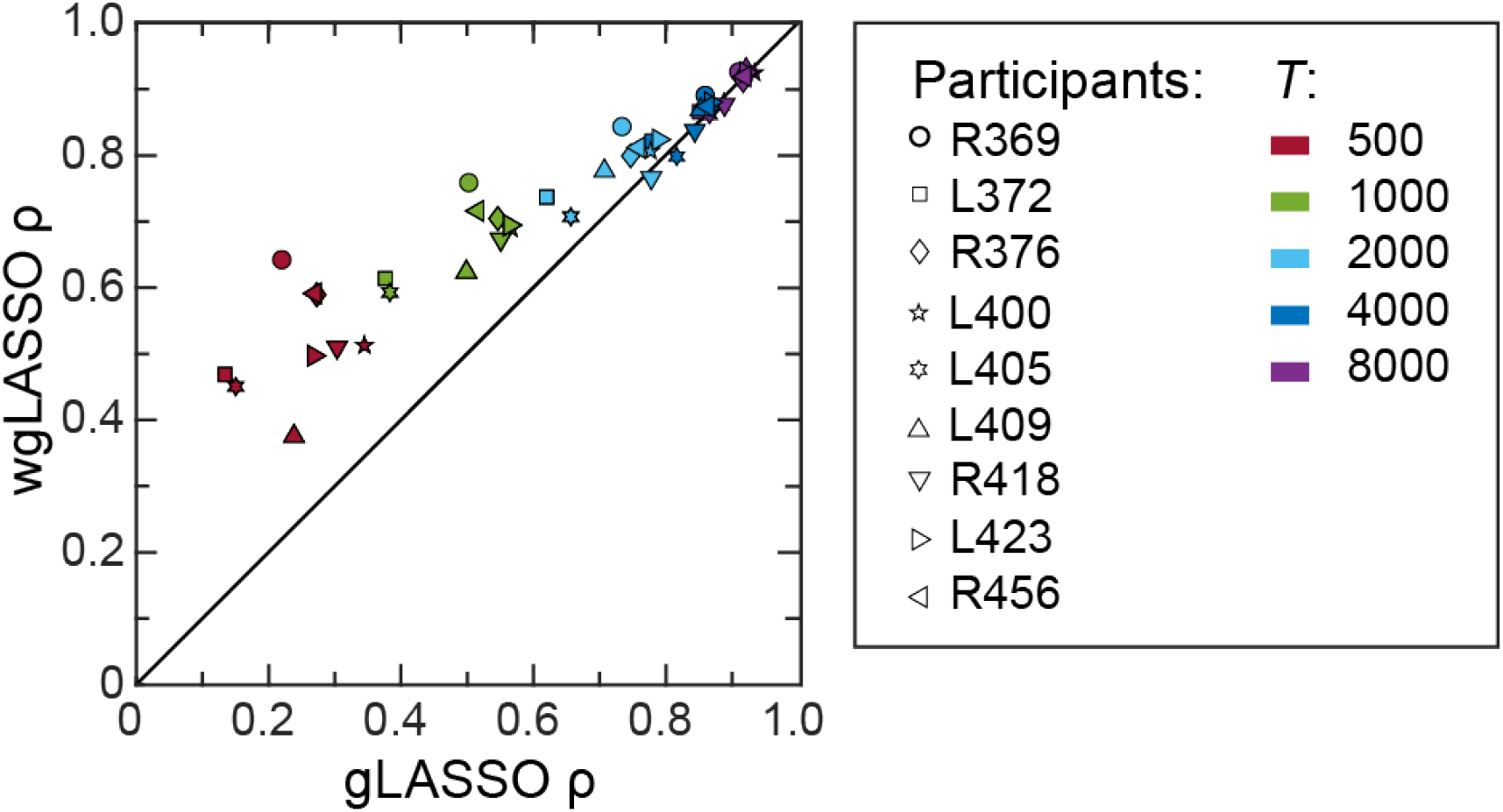
Summary of estimator performance across participants for SSGC. Cosine similarity computed between the estimated and ground truth SSGC connectivity matrices for wGLASSO plotted versus gLASSO estimators.

**Supplementary Figure 8:**
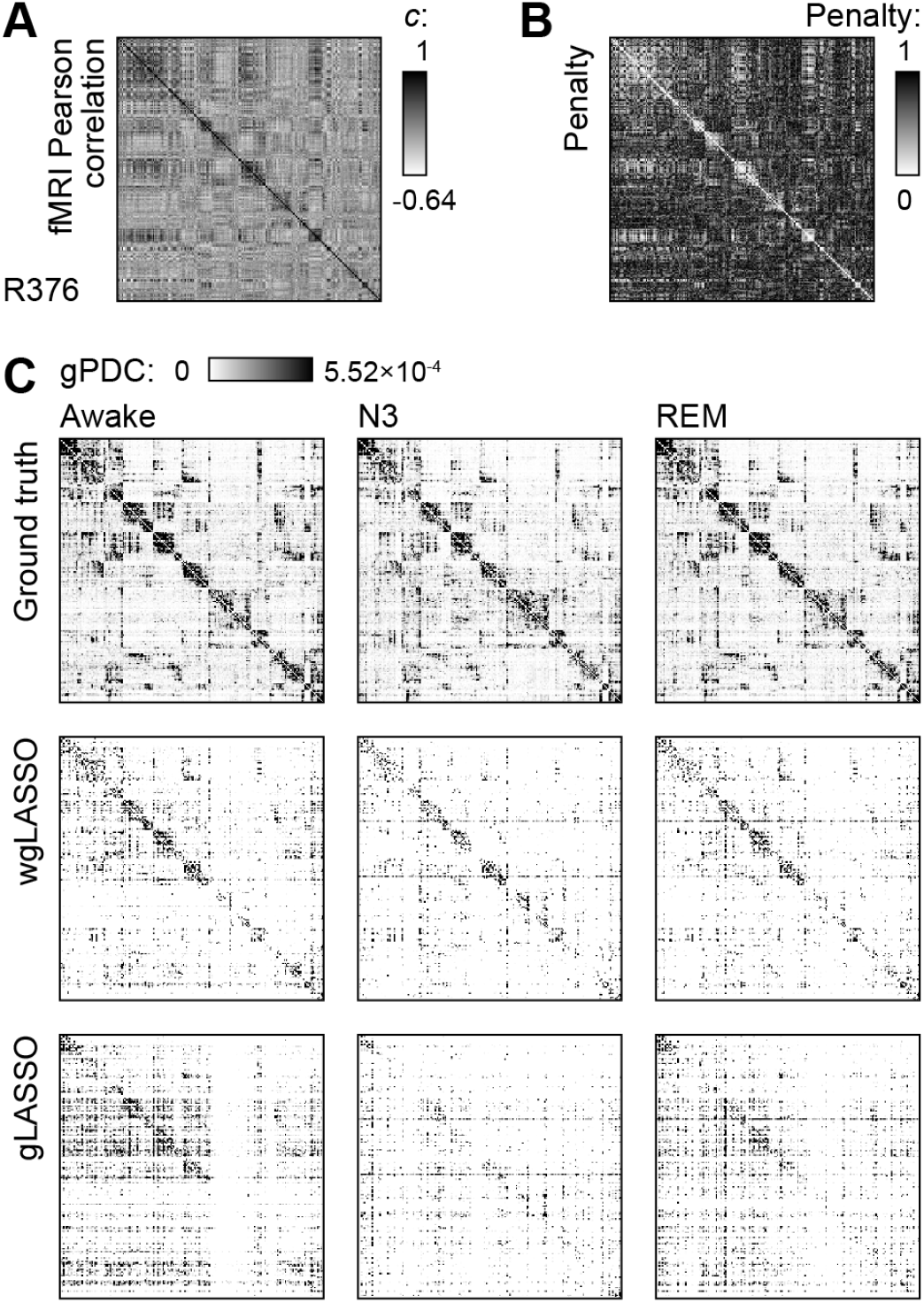
Single subject (376R) example of gLASSO and wgLASSO estimation methods applied to data recorded during sleep. **(A)**. fMRI Pearson correlation matrix for this subject, obtained during a daytime scan. **(B)**. Penalty matrix derived from the matrix in A, applied to wgLASSO estimates for all sleep stages. **(C)**. gPDC matrices derived from ground truth models (top row), wgLASSO estimates (middle row), and gLASSO estimates (bottom row) for data recorded during three stages of overnight sleep: Awake (left column), N3 (middle column), and REM (right column). Each estimated gPDC matrix is averaged across five trials for each method; 1000 training samples were used to fit the estimated models.

## Acknowledgements

This research was performed using the computer resources and assistance of the UW-Madison Center for High Throughput Computing (CHTC) in the Department of Computer Sciences. The CHTC is supported by UW-Madison, the Advanced Computing Initiative, the Wisconsin Alumni Research Foundation, the Wisconsin Institutes for Discovery, and the National Science Foundation, and is an active member of the OSG Consortium, which is supported by the National Science Foundation and the U.S. Department of Energy’s Office of Science. This research was done using resources provided by the Open Science Grid (Pordes et al., 2007), which is supported by the National Science Foundation award #2030508. Funding was provided by National Institutes of Health (grant numbers R01-DC04290, R01-GM109086, UL1-RR024979) and the UW Department of Anesthesiology. The authors are grateful to Graham Berger, Haiming Chen, Christpher Garcia, Phillip Gander, Matthew Howard, Hiroto Kawasaki, Christopher Kovach, and Ariane Rhone for help with data collection and analysis.

